# Therapeutic Effects of An Insulin-Like Growth Factor I Sensitizer In Traumatic Brain Injury

**DOI:** 10.64898/2026.05.13.724506

**Authors:** J.A. Zegarra-Valdivia, M.Z. Khan, A. Putzolu, J. Pignatelli, I. Torres Aleman

## Abstract

Traumatic brain injury (TBI) is a condition of high incidence worldwide, but remains mostly undertreated. Previous observations in preclinical studies pointed to a beneficial effect of insulin-like growth factor 1 (IGF-1) in TBI. As brain injury is associated to loss of IGF-1 sensitivity, we tested the therapeutic potential of AIK3a305 (AIK3), a novel IGF-1 sensitizer. Twenty-four hours after mild TBI induced by controlled impact, mice received daily intraperitoneal injections of AIK3 during 4 weeks. We found that TBI-associated sensorimotor disturbances measured with the adhesive-removal test were reverted by AIK3 treatment. In addition, neurological and cognitive disturbances measured by the neurological severity score and Y maze respectively, were also ameliorated by treatment with the IGF-1 sensitizer, whereas increased anxiety after mild TBI was also normalized by AIK3. Circulating levels of IGF-1 were increased after AIK3 treatment in TBI mice, while serum IL-6 levels, a biomarker of inflammation associated to TBI were similar to control mice treated with AIK3. Transcriptomic analysis determined that treatment with AIK3 widely affected gene expression in TBI brains, showing a general reduction in both up- and down-regulated genes. Collectively, these data support the use of IGF-1 sensitizers such as AIK3 for treatment of TBI.

## Introduction

The prevalence of traumatic brain injury (TBI) is among the highest of all neurological conditions worldwide (Maas et al., 2022), with up to 60 million people affected each year (Cieri and Ramos, 2025). While mild forms are the most common (Maas et al., 2022), they are associated to many invalidating morbidities (mood and cognitive disorders, epilepsy, ..etc), making the long-term impact of TBI even more detrimental (Masel and DeWitt, 2010). Although the search for new therapeutic approaches is ongoing, still no effective treatments are available, and none oriented to targeting disease mechanisms (Maas et al., 2022). Heterogeneity of neuropathology (Jassam et al., 2017), and lack of translation of pre-clinical findings into clinical practice are invoked to explain the paucity of therapies.

Insulin-like growth factor I (IGF-1) is widely neuroprotective (Fernandez and Torres-Aleman, 2012) and, as in many other brain pathologies, its local levels are upregulated in TBI (Madathil et al., 2010). Moreover, serum IGF-1 levels correlate with cognition in TBI patients (Ou et al., 2023) and its administration promotes recovery in pre-clinical models of moderate TBI (Saatman et al., 1997; Carlson and Saatman, 2018; Venkatasamy et al., 2025). Since inflammation, excitotoxicity, and oxidative stress associated to TBI (Kaur and Sharma, 2018; Kalra et al., 2022), elicit IGF-1 resistance (Venters et al., 1999; Garcia-Galloway et al., 2003; Davila and Torres-Aleman, 2008), in the present study we used AIK3a305 (AIK3) to determine its potential therapeutic utility in TBI. This compound is a recently developed IGF-1 sensitizer able to cross the blood-brain-barrier and shows beneficial actions on brain function (Zegarra-Valdivia et al., 2022b). After mild TBI, mice systemically treated with AIK3 showed marked functional recovery, suggesting that enhancing the sensitivity to endogenous IGF-1 may be sufficient to ameliorate TBI outcome.

## Materials and Methods

### Animals

Mice were used according to ARRIVE guidelines as indicated before (Zegarra-Valdivia et al., 2022a). Mice were housed in standard cages (48 × 26 cm ^2^) with 5 mice per cage. Mice were maintained on a light-dark cycle (12-12 h, lights on at 8 am) at constant temperature (22°C) and humidity, and with food (pellet rodent diet) and water *ad libitum*. All experimental protocols were performed during the light cycle and followed European guidelines (86/609/EEC & 2003/65/EC, European Council Directives). Studies were approved by the respective local Bioethics Committees UPV M20-2022-333). Animals were not randomized and were used in a sex-balanced manner throughout. Potential confounders were not accounted for. Each experimenter took account of group allocation under study. All efforts were done to reduce harm to the animals. Mice were handled for three days prior to any experimental manipulations and familiarized with behavioral arenas to minimize novelty stress or deeply anesthesized with pentobarbital prior to sacrifice, when needed. Sample sizes were kept as little as possible to comply with current animal reduction policies. No adverse events were found. End-point measures included checking reflexes in deeply anesthesized animals prior to culling.

### Materials

We used a novel IGF-1 sensitizer, AIK3a305 (Allinky Biopharma, Spain), as described before (Zegarra-Valdivia et al., 2022b). AIK3 or the vehicle (DMSO + saline) were injected intra-peritoneally (ip) at a dose of 20 mg/kg/day, 1 day after traumatic brain injury, and during 4 weeks (Suppl Figure 1A). This compound shows good blood-brain-barrier penetration and enhances responses to IGF-1 (Zegarra-Valdivia et al., 2022b).

### Traumatic brain injury (TBI)

Mild TBI was induced as described (Santi et al., 2018) using an electromagnetic stereotaxic impactor (Impact One™, MyNeuroLab, Leica, Germany). This is the most common type of TBI in the clinic. In brief, mice were anesthetized with isofluorane (induction at 4% and maintenance at 2%) and placed on a digital stereotaxic frame (Stoelting Co, USA) with cup head holders (David Kopf Instruments, USA). Temperature was maintained with a custom-made heating bed. The skin was scrubbed with Betadine^®^, and the top of the skull was exposed with a 1 cm skin incision. A craniotomy was performed using a 5 mm trephine (Fine Science Tools, USA). The right edge of the impactor tip (3 mm) was aligned with the midline suture and the posterior edge of the tip with the horizontal anterior portion of the bregma suture using the stereotaxic arm. The tip was positioned over the left parieto-temporal cortex (-1.0 mm bregma and 2.5 mm left to midline). The zero-depth position was determined by aligning the tip of the impact device in the down position with the surface of the dura. When contact was made, the impactor tip was retracted and lowered 1 mm before inducing impact (speed: 3.6 m/s dwell: 100 ms). The surface of the dura was cleaned and dried with room-temperature sterile saline and dry cotton. A 6 mm-diameter plastic disc was glued with cyanoacrylate to the skull to cover the craniotomy. The skin was closed with 4 discontinuous sterile 5-0 braided silk sutures (Lorca Marin, Spain).

### Experimental design

As shown in Figure 1A, treatment with AIK3a303 (ip) or the vehicle (DMSO 1% in saline) was initiated 24 hours after TBI. Functional and behavioral testing was conducted along 3 weeks. At the end of week 4, animals were sacrificed and blood and brain tissue collected and stored frozen until use.

**Figure 1.**
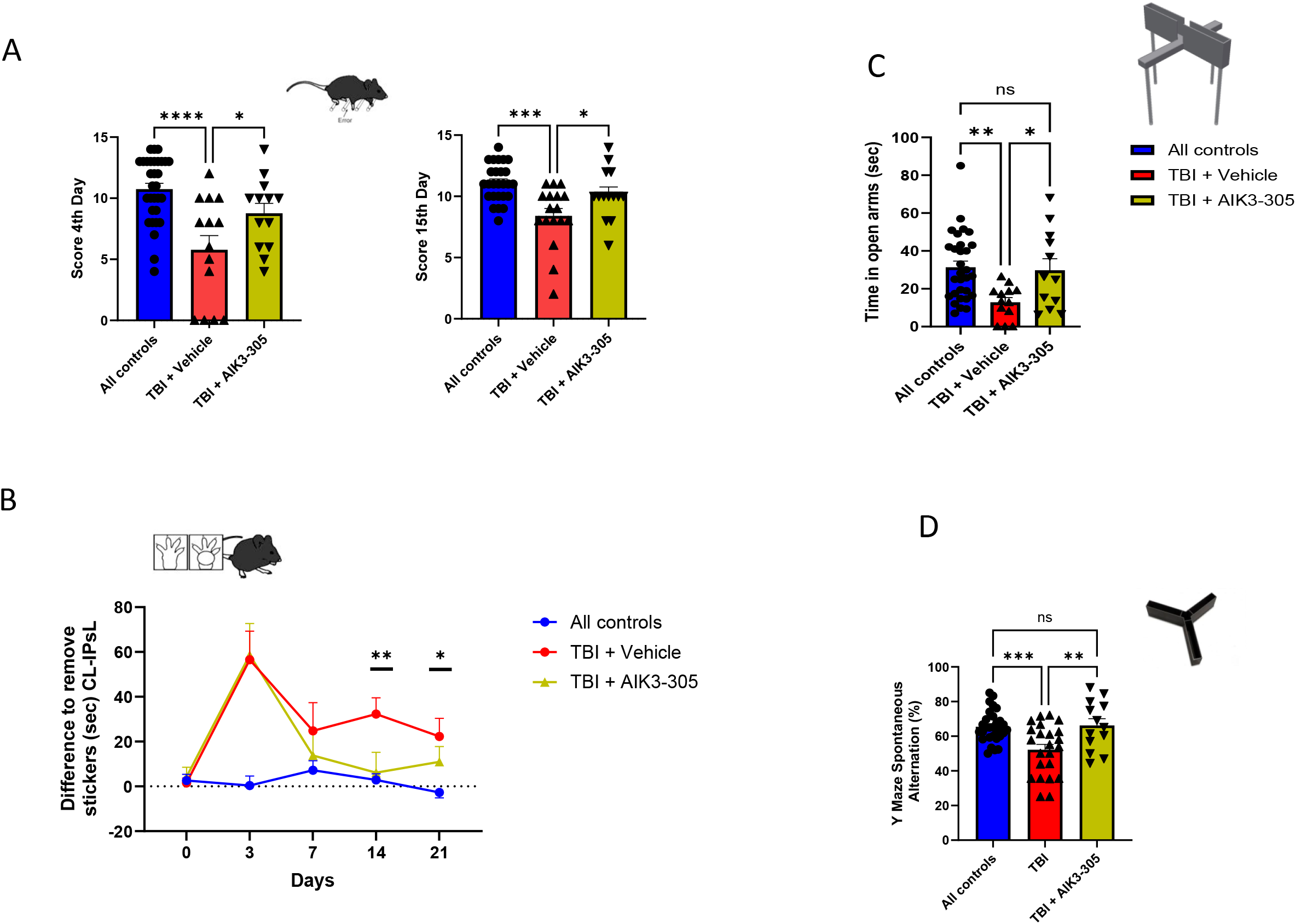
Neurological function after TBI is protected by treatment with AIK3a305. **A**, Using the neurological severity scale, it is observed that at 4 and 15 days after injury, the animals treated with AIK3a305 show no functional impairments. The animals treated with the vehicle show moderate dysfunction, as expected (n= 15). **B**, Sensorimotor coordination determined by the adhesion removal test over time after injury shows that the animals treated with AIK3a305 recover function within 1 week, while vehicle-treated mice did not fully recover function even after 4 weeks (n= 15). **C**, Increased anxiety caused by the injury is not observed in mice treated with AIK3a305, as reflected in the longer time spent in the open arms of the elevated maze test compared to vehicle-treated TBI mice (n= 15). **D**, Working memory determined in the Y maze was intact after 3 weeks of AIK3 treatment (n=15). *p<0.05, **p<0.01 by Student’s t-test after ANOVA in this and following figures.

### Adhesive Removal Test

This test was used to assess long-term sensorimotor deficits, as described (Santi et al., 2017). During the first week, mice were habituated to the experimental room for 1 h and placed 120 s in the testing box for habituation. After this acclimation period, animal cages were taken to the behavioral room 1 hour before starting the experiment. Subsequently, two adhesive tape strips (0.3 cm x 0.4 cm) were applied on each paw, switching the placement order (right or left) randomly within each group and mice were introduced in the testing box until they had removed both tape strips, for a maximum duration of 120 s. This procedure was repeated until there was no asymmetry on the time to remove the adhesive from each animal forepaw’s, and no significant differences between experimental groups were found. When all animals reached a *plateau*, they underwent surgery, and were re-tested at the adhesive removal test once weekly for a 28 days period. A camera placed under the testing box allowed video-recording of each trial. Observations were carried out by an investigator blind to the experimental conditions. In order to analyze the magnitude of asymmetry, time-to-remove the tape from the ipsilateral limb was subtracted from the time-to-remove the tape from the contralateral limb.

### Behavioral tests

#### Elevated plus maze

To assess anxiety-like/coping behavior, mice were introduced in a maze of 40 cm from the floor with two opposing arms. Two protected (closed) arms (30 cm (length) × 5 cm (wide) × 15.25 (height), and two opposing unprotected (open) arms (30 cm (length) × 5 cm (wide). Each animal was introduced in the middle of the apparatus for 5 minutes. Stress was scored as time spent in the closed arms while coping behavior was estimated by time spent in the open arms. All measures were recorded (Video Tracking Plus Maze Mouse; Med Associates, USA), and analyzed as described (Munive et al., 2019).

#### Neurological score

To determine general neurological function, a modified neurological severity scoring was performed 1 day after lesion, as described in detail elsewhere (Suda et al., 2024). Motor, sensory, and reflex responses were graded on a scale of 0 (maximal deficit) to 14 (normal).

#### Y maze

This test measures spontaneous alternation as an index of working memory (Sarter et al., 1988) and was used as described (Zegarra-Valdivia et al., 2022a). In brief, the mouse is placed at the end of one arm to move freely from side to side of the maze during an 8-min session. Videos recorded the sequence of entries and were analyzed off-line. Entrance to each arm is scored when the mouse places the hind paws entirely in the zone. Alternation was defined as successive entries into the three arms on overlapping triplet sets. Consecutive triplets were analyzed, and alternate behavior was calculated as the percentage of actual alternation (number of triplets with non-repeated entries) versus total alternation opportunities (total number of triplets), as described (Recinto et al., 2012).

### ELISA

IGF-1 and IL-6 in serum were determined using commercial immunoassays (Biotechne R&D Systems, USA) following the manufacturer’s instructions. IL-6 values below the detection limit of the assay were assigned the lowest detectable concentration of the assay for statistical purposes. For IGF-1, control and control + AIK3 groups were pooled together as they show similar values, while for IL-6 they are shown separate as AIK3 increased them in control mice.

### RNAseq

RNA was isolated from injured cortex regions or equivalent for control samples, using Qiagen RNA isolation Mini kit. Total RNA concentration was assessed by Qbit fluorimetry and RNA QC was performed using vertical electrophoresis (Biooptic, Taiwan) in an NS1 column, and Rin quality was determined by an 18/28S ratio. Only samples with RIN values superior to 7 were used. For mRNA sequencing, 1ug of total RNA was used for mRNA isolation using Illumina Poly(A) capture kit and reverse transcribed using Illumina cDNA synthesis kit. mRNA Libraries were built using Illumina RNA prep, index anchors were added to libraries and finally, each sample was indexed with specific i5 and i7 Illumina indexes. Libraries QC was assessed by vertical electrophoresis in an S3 column (Biooptic), quantified by fluorimetry (Qbit, Thermo-Fisher, USA) and libraries were normalized to 2nM in an RSB buffer.

Sequencing was performed in a NExtSeq2000 sequencer (Illumina, USA) using a P3 200 cycles flow cell. Libraries were pooled to a 750nM final loading concentration.

### Statistics

Statistical analysis was performed using GraphPad Prism 6 software (San Diego, CA, USA) and R Package (Vienna, Austria). Depending on the number of independent variables, normally distributed data (Kolmogorov-Smirnov normality test), and the experimental groups compared, we used either Student’s t-test, two-way ANOVAs, or Two-way repeated measure ANOVA, followed by Sidak’s multiple comparison test. For non-normally distributed data, we used the Mann-Whitney U test to compare two groups, Kruskal-Wallis or Friedman test, with Dunn’s multiple comparisons as a Post Hoc analysis as Scheirer-Ray Test, a non-parametric alternative to multi-factorial ANOVA. The sample size for each experiment was chosen based on previous experience and aimed to detect at least a p<0.05 in the different tests applied, considering a reduced use of animals. Results are shown as mean ± standard error (SEM) and *p* values coded as follows: *p< 0.05, **p< 0.01, ***p< 0.001. Animals were included in each experimental group randomly by the researcher. No animals were excluded from the analyses.

## Results

Moderate TBI induces early signs of neurological impairment, commonly assessed with the neurological severity score. Administration of AIK3 24h after TBI (Suppl Fig 1A) resulted in preserved neurological function, even in the early stages after the injury, as indicated by nearly normal neurological index scores at days 4 and 15 post-TBI (Figure 1A). Treatment with AIK3 significantly accelerated the recovery of sensorimotor function in TBI mice over the following weeks, as measured by the adhesive removal test conducted at weekly intervals post-injury (Figure 1B). Similarly, after 2 weeks, AIK3 normalized anxiety levels that were elevated by TBI, as determined by the EPM test (Figure 1C), as well as time spent in the center of the open field test (Suppl Fig 1B), whereas at week 3, working memory in TBI mice treated with AIK3 was normal, as determined in the Y maze (Figure 1D).

Since serum IGF-1 levels decrease during the first weeks after TBI (Santi et al., 2017; Corne et al., 2021), we measured them at week 4 and found that AIK3 significantly increased them, although at this time, TBI mice treated with the vehicle did not show a significant reduction in serum IGF-1 (Figure 2A). Since IGF-1 administration to vehicle injected control mice did not elicit changes in serum IGF-1 we pooled both control groups together (all controls). Additionally, we determined whether AIK3 treatment affects post-TBI inflammation by measuring IL-6 levels in serum, and found that after 4 weeks, IL-6 were only moderately increased after TBI, and treatment with AIK3 resulted in similar levels to both groups of control mice. Again, intact and IGF-1 treated controls were pooled together since IL-6 were similar in both groups (all controls). None of the changes seen in serum IL-6 were statistically significant.

**Figure 2.**
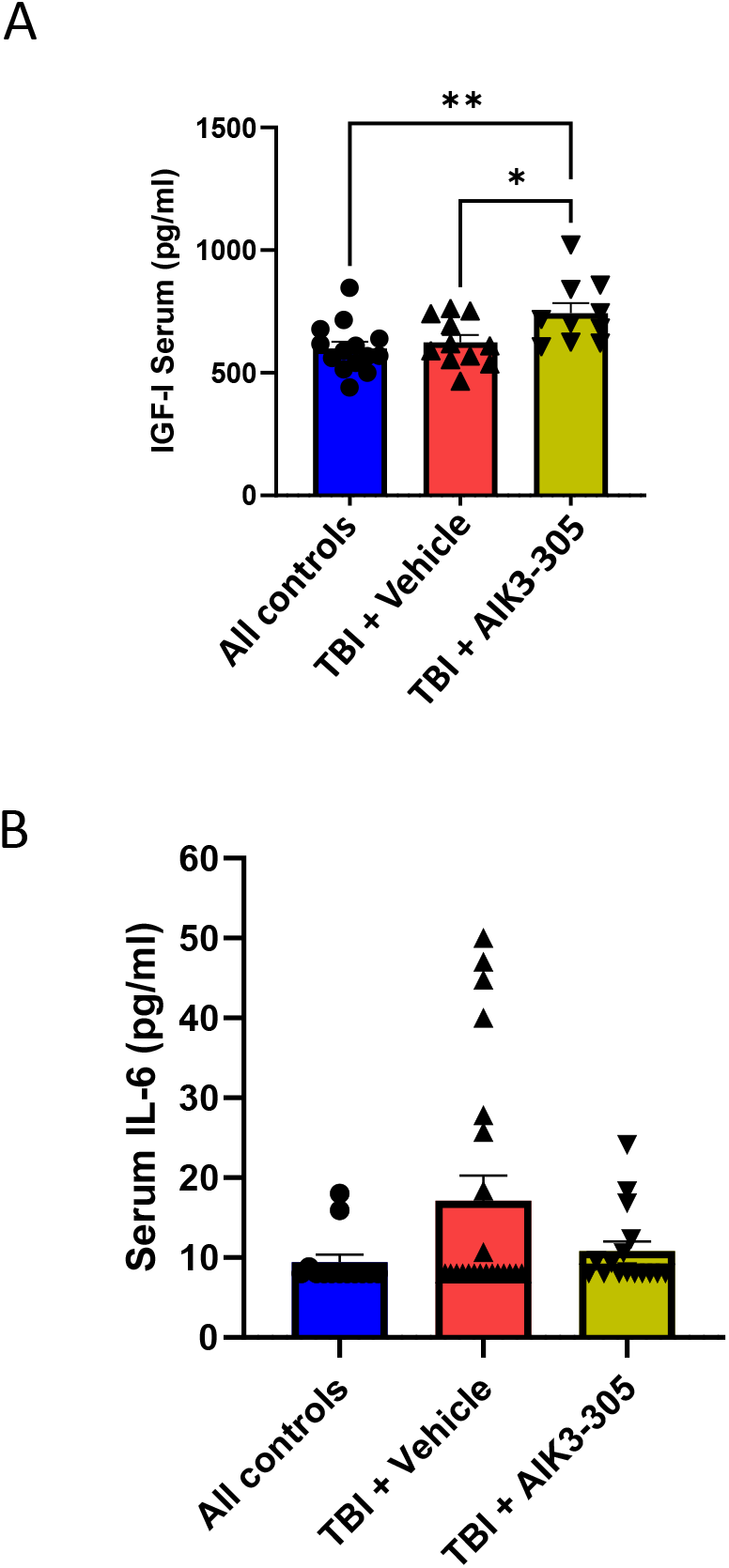
Improved TBI-associated biomarkers by AIK3 treatment. **A**, Serum IGF-1 levels were significantly increased by AIK3 in TBI mice treated for 4 weeks (n=8-10 per group). **B**, Serum IL-6 levels were not significantly increased 4 weeks after TBI, while AIK3-treated TBI mice show normal levels.

Finally, we determined possible transcriptomic changes in TBI mice treated with AIK3 and found that the number of up- and down-regulated genes compared to TBI + vehicle was reduced to a third (Figure 3A), suggesting a trend to preserve normal gene activity, although heatmap representation of changes in gene expression indicated that AIK3 treatment elicited an intermediate landscape between control and TBI groups (Figure 3B). Using a pair-wise comparison in Volcano plots, a relatively larger increase in down-regulated genes was seen after AIK3 treatment of TBI mice as compared to TBI+vehicle mice (Figure 3C). Moreover, several genes, such as CD11 (*Itgax*), cystatin F (*Cst7*) or C3, involved in phagocytosis, inhibition of proteases, or the innate immune response, respectively, were kept up-regulated after AIK3 treatment, suggesting that reparative responses were maintained (Table 1). Conversely, very few down-regulated genes after TBI remained lowered after AIK3 treatment while downregulated genes such as igfr1, that codifies for the IGF-1 receptor, Mybr, that includes transcription factors of wide function, including cell death and survival, or pak5, coding for p21-activated kinase 5, that intervenes also in cell death, were rescued by AIK3 treatment. Since the number of genes downregulated by AIK3 compared to vehicle-treated TBI mice was large, we provide a list showing the largest downregulation in Table 2.

**Figure 3.**
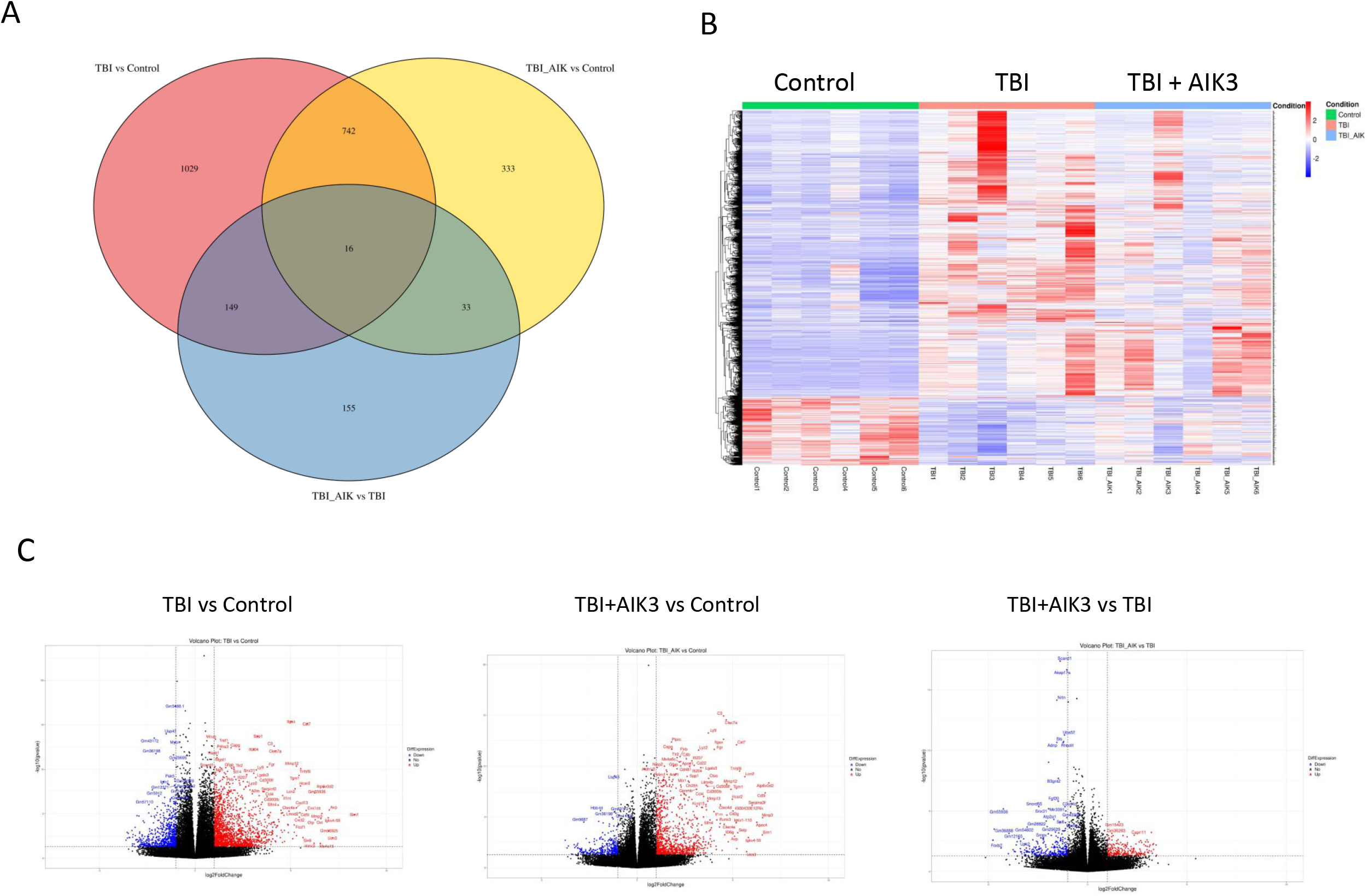
Transcriptomic analysis of AIK3 treatment in TBI mice. A, VennPlot analysis of changes in gene expression in the 3 experimental groups show over 1,000 genes changed by TBI that were reduced by AIK3 treatment to 333 genes. **B**, Heatmap representation of changes in gene expression indicates that AIK3 treatment elicited an intermediate landscape between control and TBI. **C**, Volcano plots in pair-wise comparisons again indicate that AIK3 treatment elicits a markedly different profile of gene changes.

**Table 1.** Representative genes induced by TBI that are involved in reparative responses and maintained after AIK3 administration.

| **Gene** | **Reparative response\*** |
| --- | --- |
| CD11 | Phagocytosis |
| C3 | Innate immune response |
| Cystatin F | Protease inhibition |
| MMP12 | Tissue remodeling |
| MMP3 | Wound healing |
| CD30L | Modulation of inflammatory signaling |
| Vasopressin | Vasoconstriction |
| HCAR2 | Anti-inflammation |
| Transglutaminase 1 | Cross-linking of protein networks |
| Lipocalin 2 | Modulation of cellular stress |
| Galectin 3 | Tissue remodeling |
| ATP6v0d2 | pH regulation |
| CCL4 | Recruitment of immune cells |

\*Only potential beneficial actions of these genes are included

**Table 2.**
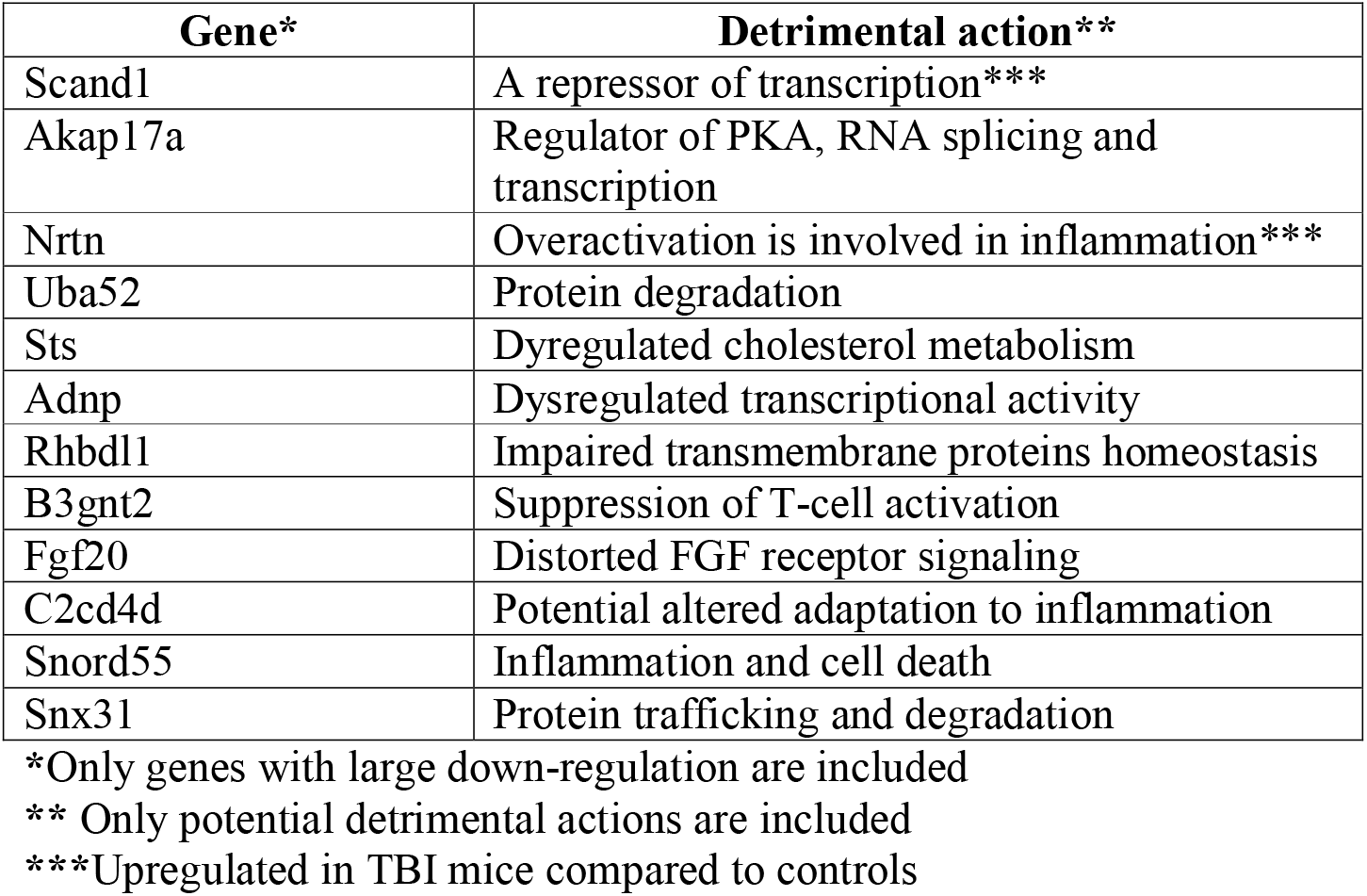
Down-regulated genes with potential deleterious actions in AIK3-treated TBI mice compared to vehicle treated.

## Discussion

These results indicate that treatment with an IGF-1 sensitizer ameliorates functional impairments produced by moderate TBI. Hence, sensorimotor disturbances produced by TBI were normalized significantly faster by AIK3 than the vehicle-treated mice, while neurological and cognitive disturbances were fully reversed by treatment with the IGF-1 sensitizer. Further, increased anxiety after mild TBI was also normalized by treatment. While we previously found that circulating levels of IGF-1 decrease early after TBI (Santi et al., 2017), after 4 weeks serum IGF-1 levels still remained, albeit moderately, decreased, and, significantly, AIK3 treatment increased them. Since decreased levels of IGF-1 have been associated to TBI in humans (Mishra et al., 2026), the effect of AIK3 may also be of interest for a future clinical use in human TBI. The pro-inflammatory cytokine IL-6 was moderately, but not significantly increased after TBI, and treatment with AIK3 lower it to levels seen in control mice treated with the compound, hinting to a potential anti-inflammatory action of the compound. Analysis of transcriptomic changes after AIK3 treatment suggest that a major action of the compound is to preserve beneficial vs detrimental gene activities triggered by TBI, suppressing those that probably hinder recovery.

Pilot studies of administration of IGF-1 in brain injuries and neurodegenerative diseases have often, but not always (Mitchell et al., 2002), shown beneficial effects (Lai et al., 1997; Nagano et al., 2005; Arpa et al., 2011), but no use of this growth factor has been approved for any brain condition yet. Based on prior successes with IGF-1 in animal models (Bozdagi et al., 2013) and clinical studies (Shcheglovitov et al., 2013; Pini et al., 2016; Kolevzon et al., 2022), a small IGF-1 mimetic (Trofinetide) was recently approved for treatment of several autistic spectrum disorders (Neul et al., 2023) and previously proved to be efficacious also in other neurological diseases (Grunseich et al., 2018). Since IGF-1 sensitizers have been shown of potential therapeutic use (Liu et al., 2017), AIK3 readily crosses the blood-brain-barrier (Zegarra-Valdivia et al., 2022b), and shows no toxicity (unpublished observations), we postulate the use of this novel IGF-1 sensitizer for TBI and other brain injuries associated to loss of sensitivity to IGF-1 due to inflammation, excitotoxicity, or oxidative stress, provided its use in the clinic is approved.

## Acknowledgements

Funding was provided by Allinky Biopharma. We thank the technical expertise provided by Achucarro Core Facilities, in particular the assistance of Dr. R Cipriani.

## LEGENDS TO FIGURES

**Supplementary Figure 1. A**, Time-line of the protocol used in the study indicating the various tests applied during the 4-week period of study. **B**, Time spent in the center of the arena of the open field test in the experimental groups (n=16).

