## Supplementary Figures for "Therapeutic Effects of An Insulin-Like Growth Factor I Sensitizer In Traumatic Brain Injury"

### Slide 1
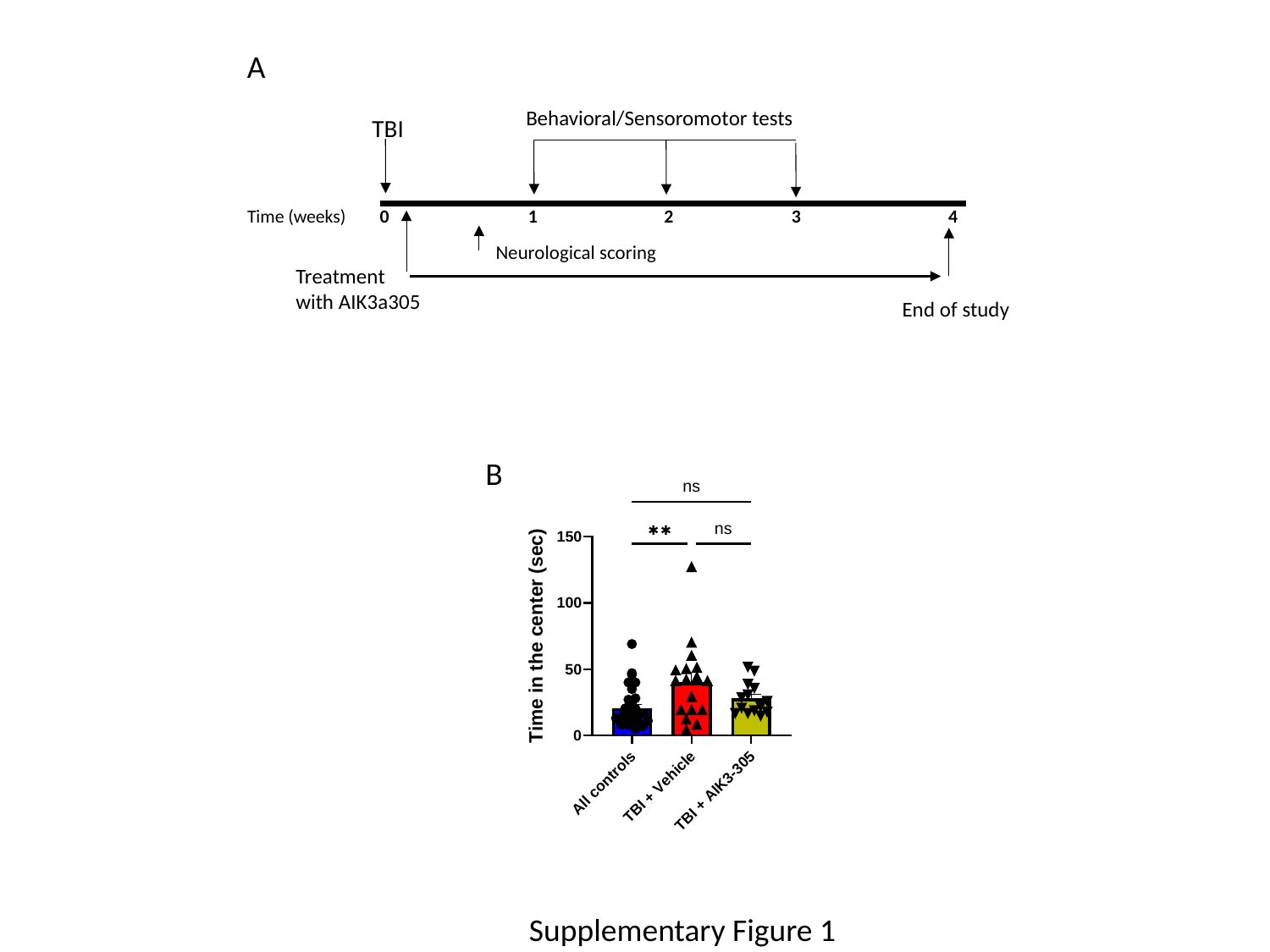

A
Behavioral/Sensoromotor tests
TBI
Time (weeks) 0 1 2 3 4
Neurological scoring
Treatment
with AIK3a305
End of study
B
Supplementary Figure 1
